# Genetic deletion of Autotaxin from CD11b^+^ cells decreases the severity of experimental autoimmune encephalomyelitis

**DOI:** 10.1101/850149

**Authors:** Ioanna Ninou, Ioanna Sevastou, Christiana Magkrioti, Eleanna Kaffe, George Stamatakis, Spyros Thivaios, George Panayotou, Junken Aoki, George Kollias, Vassilis Aidinis

**Affiliations:** Biomedical Sciences Research Center Alexander Fleming, Athens, Greece; Graduate School of Pharmaceutical Sciences, Tohoku University, Sendai, Miyagi, Japan

**Keywords:** Autotaxin (ATX), lysophosphatidic acid (LPA), experimental autoimmune encephalomyelitis (EAE), macrophages, microglia

## Abstract

Autotaxin (ATX) is a secreted lysophospholipase D catalyzing the extracellular production of lysophosphatidic acid (LPA), a growth factor-like signaling lysophospholipid. ATX and LPA signaling have been incriminated in the pathogenesis of different chronic inflammatory diseases and various types of cancer. In this report, deregulated ATX and LPA levels were detected in the spinal cord and plasma of mice during the development of experimental autoimmune encephalomyelitis (EAE). Among the different sources of ATX expression in the inflamed spinal cord, F4/80^+^CD11b^+^ cells, mostly activated macrophages and microglia, were found to express ATX, further suggesting an autocrine role for ATX/LPA in their activation, an EAE hallmark. Accordingly, ATX genetic deletion from CD11b^+^ cells attenuated the severity of EAE, thus proposing a pathogenic role for the ATX/LPA axis in neuroinflammatory disorders.

## Introduction

Multiple sclerosis (MS) is the most common debilitating disorder of the central nervous system (CNS), imposing a substantial personal, and socioeconomic burden [1]. MS is a chronic inflammatory, autoimmune and neurodegenerative disease, characterized by loss of myelin sheaths and oligodendrocytes in the brain and spinal cord, resulting in aberrant axonal conduction leading to neurologic disabilities and, eventually, impaired mobility and cognition [1]. Despite the inherent limitations of modeling a human disease in experimental animals, valuable insights into MS pathophysiology were provided by the model of experimental autoimmune encephalomyelitis (EAE), that can be induced by immunizing animals against myelin antigens [2]. In this context, current research suggests that inflammation is a prominent feature of MS pathogenesis, initiated by autoreactive lymphocytes in the periphery and substantiated by leakage of the blood-brain barrier (BBB)[3]. Neuroinflammation leads to microglial and astrocyte activation, astrogliosis and eventually to demyelination, axonal or neuronal loss [3]. Microglia and macrophages are well recognized as essential players in CNS diseases including MS, while macrophage infiltration predominates the CNS inflammatory response in EAE [4].

Autotaxin (ATX, ENPP2) is a secreted glycoprotein widely present in biological fluids, catalyzing the extracellular hydrolysis of lysophosphatidylcholine (LPC) to lysophosphatidic acid (LPA), a bioactive growth factor-like lysophospholipid [5–7]. LPA signals through at least six receptors (LPA 1-6) that exhibit widespread cell and tissue distribution and overlapping specificities [8]. LPARs couple with G-proteins, crucial molecular switches that activate many overlapping signal transduction pathways, leading in pleiotropic effects in almost all cell types, including neuronal ones [8].

ATX expression and extracellular LPA production have been shown to be essential for embryonic development; ATX deficient embryos exhibited severe neural tube defects and impaired neurite outgrowth, that could be restored by LPA *ex vivo*, indicating a major role for ATX/LPA in CNS development [9]. Moreover, ATX has additional, LPA-independent, roles in CNS development, as it was shown to modulate oligodendrocyte physiology via its matricellular properties and to affect the localization and adhesion of neuronal progenitors [10, 11].

In adult healthy life, the CNS is one of the highest ATX expressing tissues, predominately expressing the CNS specific ATX γ isoform which, in comparison to the most abundant β isoform, contains a 25 aa insert of unknown function [12, 13]. ATX has been reported to be constitutively expressed by choroid plexus and leptomeningeal cells, releasing ATX in the cerebrospinal fluid (CSF), while increased ATX levels have been detected in activated astrocytes following neurotrauma, as well as in neuroblastomas and glioblastomas [14, 15]. PLA2/ATX-dependent LPA/LPAR1 signaling has been shown crucial for the initiation of neuropathic pain [8, 16], while deregulated ATX and LPA levels have been detected, in conflicting reports, in the sera and CSF of MS patients [17–21]. Given the established role of ATX/LPA in CNS development, its likely involvement in CNS pathophysiology, as well as the reported LPA effects in different neuronal cell types [8], in this report we examined a possible role of ATX/LPA in the pathogenesis of EAE.

## Materials and Methods

### Mice

Mice were housed and bred at 20–22°C, 55±5% humidity, and a 12-h light-dark cycle at the local animal facilities under specific pathogen-free conditions; water and food were given *ad libitum*. All reported experimentation in mice for this project, in line with the ARRIVE guidelines, was approved by the Institutional Animal Ethical Committee (IAEC) of BSRC Alexander Fleming and the Veterinary service and Fishery Department of the local governmental prefecture respectively (#449 and #1369/3283/3204 respectively). The generation and genotyping protocols for *Enpp2*^f/f^ [9], *Enpp2*^f/-^ [9], Tg*Enpp2*^+/+^ [22], Tg*CD11b-Cre* [23] genetically modified mice have been described previously. All mice were bred in their respective genetic backgrounds (C57Bl6/J) for over 10 generations. All randomly assigned experimental groups consisted of littermate male age-matched mice. All measures were taken to minimize animal suffering and distress; no invasive or painful techniques were performed requiring anesthetics or analgesics. The health status of the mice was monitored at least once per day; no unexpected deaths were observed. Clinical scoring was reported as indicated in the corresponding figures. Euthanasia was humanly performed in a CO_2_ chamber with gradual filling followed by exsanguination, at predetermined timepoints.

### Experimental autoimmune encephalomyelitis (EAE)

EAE was induced in 10-12-week-old C57Bl6/J (H-2^b^) male mice following a widely used EAE protocol (Fig. 1A)[2], essentially as previously reported [24, 25]. Mice were subcutaneously immunized with 100 μg of 35-55 myelin oligodendrocyte glycoprotein (MOG_35-55_, MEVGWYRSPFSRVVHLYRNGK, GeneCust), emulsified in Freund’s Adjuvant supplemented with 1mg of heat-inactivated *Mycobacterium tuberculosis* H37RA (Difco Laboratories) to the side flanks. In addition, mice received two intraperitoneal injections of 100 ng pertussis toxin at the time of immunization and 48h later. Mice were weighed and monitored for clinical signs of EAE throughout the experiment. EAE symptoms were scored as follows: 0, no clinical disease; 1, tail weakness; 2, paraparesis (incomplete paralysis of 1 or 2 hind limbs); 3, paraplegia (complete paralysis of 1 or 2 hind limbs); 4, paraplegia with forelimb weakness or paralysis; 5, dead or moribund animal. At the day of sacrifice, blood plasma and spinal cord tissue were harvested and stored.

**Fig 1.**
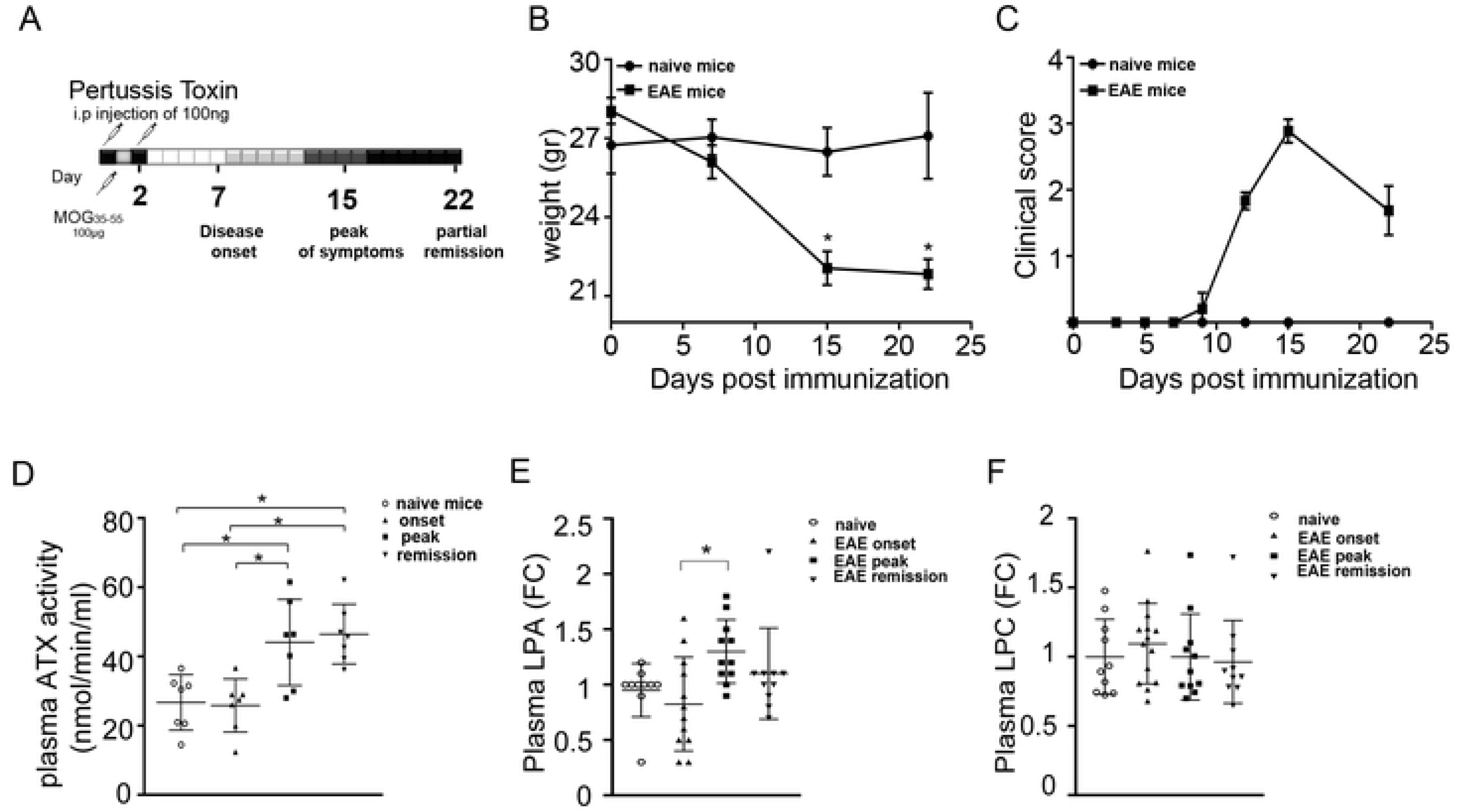
Increased ATX and LPA levels in plasma during EAE pathogenesis. (A) Schematic representation of the EAE protocol. Mice were (B) weighed and (C) monitored for clinical signs of EAE throughout the experiment. (D) Plasma ATX activity (nmol/min/ml) as measured with the TOOS assay; values are presented as mean (±SD). Total LPA (E) and LPC (F) levels in plasma; values were normalized to internal standards and presented as fold change (means ±SEM) to control samples. Statistical significance between experimental groups was assessed with one-way ANOVA complemented with Bonferroni or Dunn’s multiple pair test accordingly; * denotes statistical significance (p<0.05).

### Pathology and Immunohistochemistry

Mouse spinal cords were embedded in OCT and cryopreserved at −80°C. 7 μm sections were sliced transversely into super frost glass slides. Sections were prepared and rehydrated for Luxol fast blue staining of myelin and counterstained with hematoxylin/eosin (H&E) according to standard protocols [26].

Immunocytochemistry sections were left to dry and then were fixed in 4% paraformaldehyde for 20 min at room temperature. Sections were then permeabilized with 0.2% Triton-X for 5 min for intracellular antigen detection wherever it was necessary. Non-specific antigen sites were blocked with blocking solution (Zytomed) for 5 min, followed by addition of rabbit anti-mouse ATX (1:500, Cayman and/or Sigma) or rabbit IgG isotype control antibodies in 2% BSA at 4°C overnight. All washes were performed using PBS-Tween 0.05%. The following day the anti-rabbit Alexa555 (Abcam, 1:1000) secondary antibody was applied to the sections for 1h at room temperature, followed by counter-staining with DAPI (Fluoroshield with DAPI histology mounting medium, Sigma).

Histology images were obtained using a Nikon Eclipse E800 microscope (Nikon Corp., Shinagawa-ku, Japan) attached to a Q Imaging EXI Aqua digital camera, using the Q-Capture Pro software. Immunofluorescence images were captured under a Zeiss Axiovert200 microscope (Carl Zeiss, Oberkochen, Germany).

### Western blot

Mouse spinal cords were flushed from the spinal column, snap frozen in liquid nitrogen and stored at −80°C thereafter. Tissue was homogenised with a glass-glass homogeniser in lysis buffer containing protease inhibitors leupeptin, pepstatin and phenylmethanesulfonylfluoride. Following centrifugation at 17000 g the supernatant (cytoplasmic and soluble proteins) was collected for analysis with western blotting. Protein concentration was determined with the Bradford assay using a standard curve of BSA (0,125-2mg/ml). Proteins were separated by 8% SDS-PAGE and transferred to Protran nitrocellulose membranes (GE Healthcare, Bucks, UK) using the Trans-Blot SD Semi-Dry Transfer system (Bio-Rad Laboratories, CA, USA). Primary anti-ATX Ab incubation (monoclonal 4F1, 1:1000) was performed overnight in 2.5% (wt/vol) non-fat milk at 4°C. The membranes were then washed three times with TBS-Tween 0.05% and incubated with an anti–rat HRP-conjugated secondary Ab (1:1000) for 1 h at room temperature. Membranes were washed three times with TBS-Tween 0.05%, and antibody–antigen complexes were revealed using luminol as a chemiluminescent reagent.

### ELISA

Duplicate samples and standards in coating buffer (0,012M NaCO_3_ and 0,028M NaHCO_3_, pH 9,6) were incubated overnight at 4°C. Samples were diluted 1:100 after titration was performed. Custom made autotaxin (ENPP2-8H, AGF06181012)(Ascent Gene, MD, USA) at concentrations (100-3.125) ng/ml was used to construct a linear standard curve. After blocking for 1.5 hours with 1.5%BSA in PBS-T, samples were incubated with a-ATX rabbit anti-mouse antibody 1:1000 (10005373, Cayman, Tallinn, Estonia) for 1 hour. The a-ATX antibody was detected with an a-rabbit HRP conjugated antibody 1:2000 (4010-05, Southern Biotech, AL, USA). Colour was developed using TMB (3,3’,5,5’-Tetramethylbenzidine, A7888, Sigma, USA). The reaction was stopped with 2M H_2_SO_4_ and readings were obtained at 450nm.

### ATX activity assay

Plasma samples (100-fold diluted) were incubated in LysoPLD buffer (100 mM Tris-HCl pH 9.0, 500 mM NaCl, 5 mM MgCl_2_, 5 mM CaCl_2_, 60 μM CoCl_2_, 1 mM LPC) at 37°C for 4 hours at a final volume of 100 μl in a 96-well plate. At the end of the incubation, a colour mix (0.5 mM 4-AAP, 7.95 U/ml HRP, 0.3 mM TOOS, 2 U/ml choline oxidase in 5 mM MgCl_2_/ 50 mM Tris-HCl pH 8.0) was prepared and 100 μl were added to each well. Absorbance (A) was measured at 555 nm every 5 minutes for 20 minutes. For each sample, the absorbance was plotted against time and the slope (dA/min) was calculated for the linear (steady-state) portion of each reaction. ATX activity was calculated according to the following equation: Activity (U/ ml) = (μmol/ min/ ml) = [dA/ min (sample) - dA/ min (blank)] * Vt/ (e* Vs* 0.5) where Vt: total volume of reaction (ml), Vs: volume of sample (ml), e: milimolar extinction coefficient of quinoneimine dye under the assay conditions (e = 32,8 μmol/ cm^2^) and 0.5: the moles of quinoneimine dye produced by 1 mol of H_2_O_2_.

### HPLC MS/MS

Lipidomics was performed with an LC/ESI-MS/MS method (Thermo Scientific Dionex UltiMate 3000 RSLC / ESI-LTQ Orbitrap XL hybrid FT-MS). Lipids were extracted with a two-step recovery procedure. The first extraction (LPC extraction) recovers the most abundant lipids (PC, LPC, TG, DAG) that can be degraded artificially by the acidification and excludes them from the low abundant lipids. Lipid extraction from spinal cord and plasma was performed on ice, according to the Folch method with minor modifications. Plasma (50 ul) and homogenized spinal cord (9-74 mg) in saline were mixed with 900 μl of ice-cold PBS spiked with the internal standard mixture (17: 0 LPA, 17: 0 LPC, 12: 1 Ceramide, 17: 0 1-Sphingosine Phosphate). The samples were extracted using 2mL of ice cold CHCl_3_/CH_3_OH (2/1 v/v), followed by thorough mixing for 1min and centrifugation at 4°C for 5min at 1200×g. The lower organic layer was collected in a fresh siliconized tube and the supernatant was washed with 2 ml of PBS saturated ice cold CHCl_3_/CH_3_OH (2/1 v/v) and the lower organic layer was collected as before. The lower organic phases from both extraction steps were pooled and maintained on ice.

For the second step of the extraction (LPA extraction), the remaining aqueous phase from LPC extraction was left on ice for 10 minutes and extracted twice with ice cold CHCl_3_ / CH_3_OH (2/1, v / v) acidified (pH 3-4) with HCL. The lower organic phases were pooled and neutralized to pH 6-7 (checked with pH paper). Both organic phases from LPC and LPA extraction were evaporated to dryness and re-dissolved in 150 μL of isopropanol. After a short vortex of the solution, the sample was transferred to the HPLC sample vial for analysis loaded on a Phenomenex Luna Silica column and eluted with a binary gradient. The mobile phase was delivered at a flow rate of 0.3 mL/min and the total run time was 50 min/sample. The recovery of all lipids ranged between 60% and 100%. The relative standard deviation (RSD) was less than 7% in plasma containing three different concentrations of LPA (0.1, 0.5 and 1 μM) and of LPC (0.5, 5 and 50 μM). The analysis was performed by electrospray ionization (ESI) in negative mode for the lipids recovered from the acidified fraction (LPA) and positive for lipids derived from the neutral fraction (LPC). Data were collected and analyzed with the Xcalibur software package. The precursor ion mass was used for quantitation, while the identity of each lipid was confirmed by its fragments obtained from Collision Induced Dissociation (CID) fragmentation technique. Lipid concentration was normalized against the spiked in internal standard, in the case of lipid classes that there was no available standard, normalization was based on the closest in structure lipid standard present in the sample.

### RNA extraction and Q-RT-PCR analysis

Mouse spinal cords were flushed from the spinal column, snap frozen in liquid nitrogen and stored at −80°C thereafter. RNA was extracted from SC tissue with Tri Reagent (Molecular Research Center, OH, USA) and treated with DNAse (RQ1 RNAse-free DNAse, Promega,Wis, USA) in accordance to the manufacturer’s instructions. Reverse transcription of 3.5 μg isolated RNA was achieved with M-MLV reverse transcriptase (M1705, Promega, WI, USA) following the manufacturer’s instructions. Real-time PCR was performed on a BioRad CFX96 Touch™ Real-Time PCR Detection System (Bio-Rad Laboratories Ltd, CA, USA). Values were normalized to the expression values of hypoxanthine-guanine phosphoribosyltransferase (HPRT). The (5’-3’) sequences for forward (f) and reverse (r) primers were as follows: *Hprt* f: GGC CAG ACT TTG TTG GATTT; r: CAG ATT CAA CTT GCG CTC AT. *Enpp2 (ps1)* f: GAT GCA TTC CTT GTA ACC AAC A; r: TCA TCC TCA ATG TCA CGT AAG C. *Enpp2 (ps2)* f: GTG AAA TAT TCT TAA TGC CTC TCT G; r: GCC TTC CAC ATA CTG TTT AAT TCC. *Enpp2-γ* f: GAA ACC GGA AAA TTC AGA GG; r: CAC TTT CAA AGT CCG TAT GG. *Il-6* f: TAG TCC TTC CTA CCC CAA TTT CC; r: TTG GTC CTT AGC CAC TCC TTC. *Il-10* f: GCT CCT AGA GCT GCG GAC T; r: TGT TGT CCA GCT GGT CCT TT. *Tnf-a* f: CCT GTA GCC CAC GTC GTA G; r: GGG AGT AGA CAA GGT ACA ACC C. *Tgf-b* f: CTC CCG TGG CTT CTA GTG C; r: GCC TTA GTTT GGA CAG GAT CTG. *Col1a1* f: `CTA CTA CCG GGC CGA TGA TG; r: CGA TCC AGT ACT CTC CGC TC`. *Col3a1* f: `GCC CAC AGC CTT CTA CAC; r: CCA GGG TCA CCA TTT CTC`. *Col4a1* f: CAG GTG TGC GGT TTG TGA AG; r: TGG TGT GCA TCA CGA AGG AA. *Lpar1* f: `GAG GAA TCG GGA CAC CAT GAT; r: TGA AGG TGG CGC TCA TCT`. *Lpar2* f: `GAC CAC ACT CAG CCT AGT CAA GAC; r: CAG CAT CTC GGC AGG AAT`. *Lpar3* f: GCT CCC ATG AAG CTA ATG AAG ACA; r: TAC GAG TAG ATG ATG GGG. *Lpar4* f: AGT GCC TCC CTG TTT GTC TTC; r: GCC AGT GGC GAT TAA AGT TGT AA. *Lpar5* f: ACC CTG GAG GTG AAA GTC; r: GAC CAC CAT ATG CAA ACG. *Lpar6* f: GAT CAC TCT CTG CAT CGC TGT TTC; r: CCC TGA ACT TCA GAG AAC CTG GAG. *Pla2g1b* f: CAC CCC AGT GGA CGA CTT AG; r: GCA TTT GTT GTT TTT GGC GCT. *Pla2g3* f: AGA GAC CAC AGG GCC ATT AAG; r: GCT GTA GAA TGA CAT GGT GCT. *Pla2g6* f: GCA AGC TGA TTA CCA GGA AGG; r: GAG AGA AGA GGG GGT GAG TTG. *Pla2g12a* f: GCA ACG GCA TCC ACA AGA TAG; r: CAT AGC GTG GAA CAG GCT TC. *Pla2g4a* f: CAG CAC ATT ATA GTG GAA CAC CA; r: AGT GTC CAG CAT ATC GCC AAA. *Pla2g4c* f: TGG CTG GGA ATC CTG GGA A; r: GAG AGC ACA GGT GGT GAG TC. *Pla2g4E* f: ATG GTG ACA GAC TCC TTC GAG; r: CCT CTG CGT AAA GCT GTG G. *Pla2g4f* f: AGC CAT ACT GCT ACG GAA GAC; r: TTT GGA CAA CTT ATC TGT GTG CT. *Pla2g16* f: GGA CCC AAG CAA AGG CAT CC; r: CCA GCT CCT GCG ATT TCA CT. *Pla2g7* f: CTT TTC ACT GGC AAG ACA CAT CT; r: CGA CGG GGT ACG ATC CAT TTC. *Plpp1* f: TGT ACT GCA TGC TGT TTG TCG CAC; r: TGA CGT CAC TCC AGT GGT GTT TGT. *Plpp2* f: TCC TTT GGC ATG TAT TGC ATG T; r: AAG GCC ACC AAG AAG AAC TGA. *Plpp3* f: ATA AAC GAT GCT GTG CTC TGT GCG; r: TTT GCT GTC TTC TCC TCT GCA CCT.

### Flow cytometry

Mononuclear cell suspensions of spinal cords were prepared according to a standard protocol [27]. Briefly, spinal cord tissue was homogenized with 1X HBSS in the presence of Golgi Plug (1:1000, BD Biosciences, San Jose, CA) and passed through nylon mesh filter for further dissociation. Monocytes from SCs were isolated from the interphase of a gradient isotonic Percoll (30/70%, Sigma) at room temperature. Singlecell suspensions from spinal cord were incubated with Fc block (CD16/32-clone 93, BioLegend) and then stained with fluorescent-conjugated antibodies for surface markers: Alexa700 anti-mouse CD45 (BioLegend), APC-Cy7 anti-mouse CD11b (BioLegend) and phycoerythrin (PE) anti-mouse F4/80 (BD Biosciences). For intracellular staining, cells were permeabilized (Permeabilization wash buffer 10X, BioLegend) and stained with rabbit anti-mouse ATX (Cayman), followed by incubation with a FITC-conjugated anti-rabbit antibody. The appropriate isotype control (rabbit IgG) was used. Data was collected on a FACsCalibur and analyzed by FlowJo software (Tree Star, Ashland, OR).

### Statistical analysis

In EAE, statistical significance at specific timepoints between experimental groups was assessed with Mann-Whitney Rank Sum Test using SigmaPlot 11.0 (Systat Software, IL, USA). Otherwise, following confirmation of a normal distribution, one-way ANOVA with post hoc correction or student t-test were used accordingly and as indicated in each figure legend. Values are presented as means (±SEM). * indicates a statistical significance difference between the indicated groups (p <0.05).

## Results

### Increased ATX activity and LPA levels in the plasma upon EAE

To explore a possible involvement of the ATX/LPA axis in the pathophysiology of EAE, C57Bl6/J male mice were immunized with myelin oligodendrocyte glycoprotein (MOG), following a widely used EAE protocol (Fig. 1A)[2]. Immunized mice were weighted (Fig. 1B) and monitored macroscopically daily for clinical signs of EAE for 22 days, in comparison with naïve littermate mice, as shown for the onset (d7), peak (d15) and remission (d22) timepoints (Fig.1 A, C), where blood plasma and spinal cord tissue samples were collected.

Increased ATX activity levels were detected in the plasma of mice at the peak of EAE clinical symptoms with a well-established enzymatic assay (TOOS; Fig. 1D). To correlate ATX expression levels with its enzymatic substrate and product, we then performed HPLC/MS/MS lipidomic analysis in the plasma. Plasma total LPA levels were also found increased (Fig. 1E and S1A). No statistically significant deregulation of LPC total plasma levels was recorded (Fig. 1F and S1A), while no direct correlation of LPC with LPA species was observed (Fig. S1), as is also the case for healthy conditions [5].

To examine if the increased ATX/LPA levels in the circulation are sufficient to modulate EAE pathogenesis, EAE was induced in the heterozygous complete knock out mouse for ATX (*Enpp2*^+/−^), that presents with 50% of normal serum ATX/LPA levels [9], as well in homozygous transgenic mice overexpressing ATX in the liver (Tg*Enpp2*^+/+^), driven by the human α1-antitrypsin inhibitor (*a1t1*) promoter, resulting to 200% normal serum ATX/LPA levels [22]. No differences were observed between *Enpp2*^+/−^ mice (Fig. 2 A-C) or Tg*Enpp2*^+/+^ mice (Fig. 2 D-F) and their wild type, littermate controls in EAE incidence, onset or severity. Therefore, systemic (but lifelong) 2-fold fluctuations of ATX/LPA levels *per se*, are not enough to modulate pathogenesis in this EAE model.

**Fig 2.**
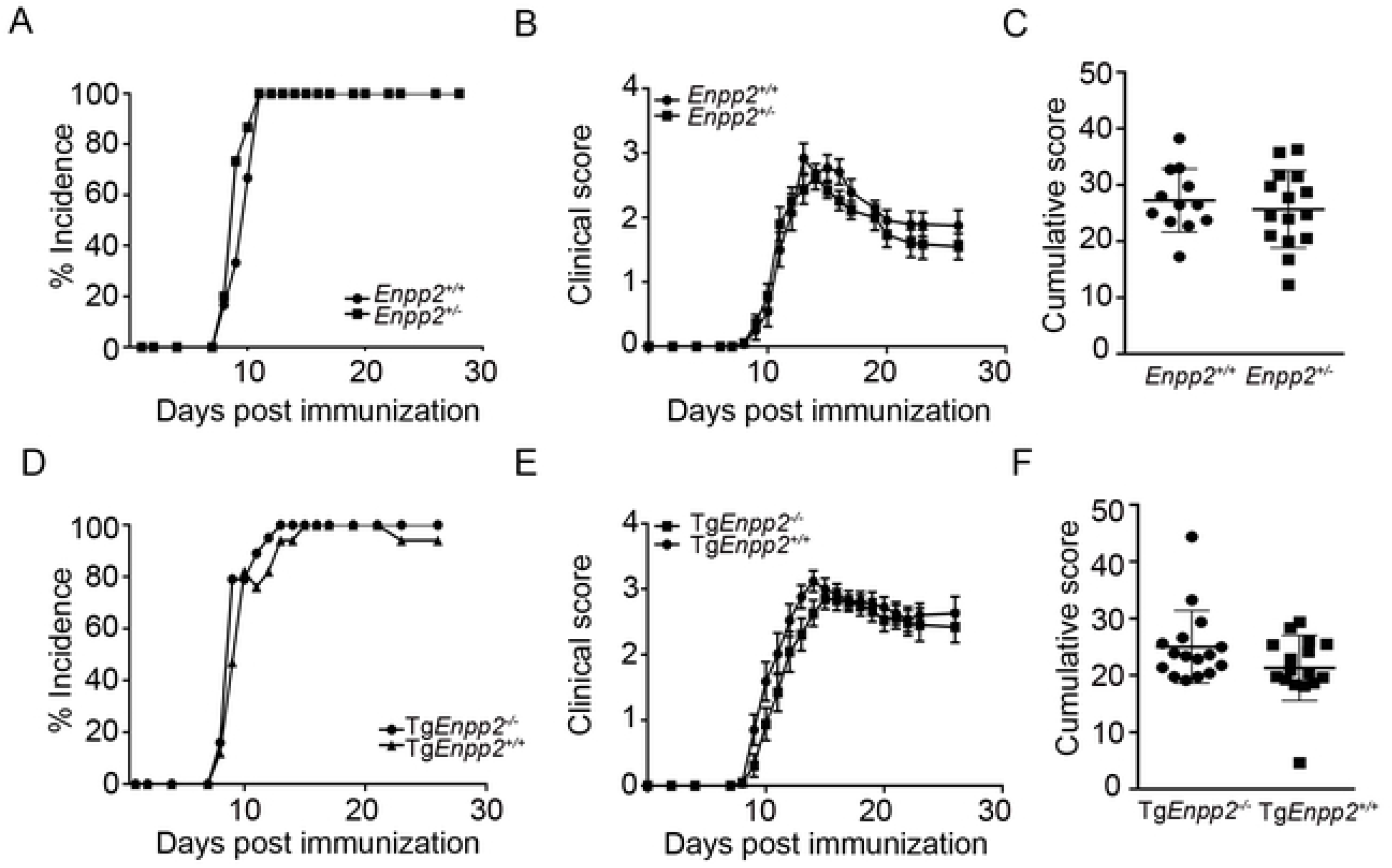
Systemic 2-fold fluctuations of ATX levels do not affect EAE pathogenesis. (A, D) EAE incidence in *Enpp2*^+/−^ and Tg*Enpp2*^+/+^ mice respectively, in comparison with the corresponding littermate controls. (B, E) Clinical and (C, F) cumulative scores of EAE progression in *Enpp2*^+/−^ and Tg*Enpp2*^−/−^ mice respectively. Values are presented as mean (± SEM). Statistical significance between experimental groups was assessed with Mann-Whitney Rank Sum Test (B and E) or Student t-test (C and F); *p<0.05.

### Increased ATX levels and deregulated lipid homeostasis in the inflamed spinal cord

In order to examine if LPA homeostasis and signaling are perturbed upon EAE locally in spinal cords, where EAE predominately manifests in this model [2], we next examined ATX and LPA levels, as well as the mRNA levels of several key players in LPA metabolism and signaling.

Increased ATX (α-ε) mRNA levels were detected with Q-RT-PCR (using two different sets of primers amplifying different parts of the mRNA) in the spinal cords of mice throughout the development of EAE (Fig. 3A), peaking at the onset and remission phases; increased mRNA levels were also detected for the CNS-specific ATX γ isoform at the same disease’s timepoints (Fig 3A). No direct correlation of ATX mRNA expression profile was apparent with the mRNA expression of inflammatory or fibrotic genes (Fig. S2 A-C), that could possibly explain the observed expression profile.

**Fig 3.**
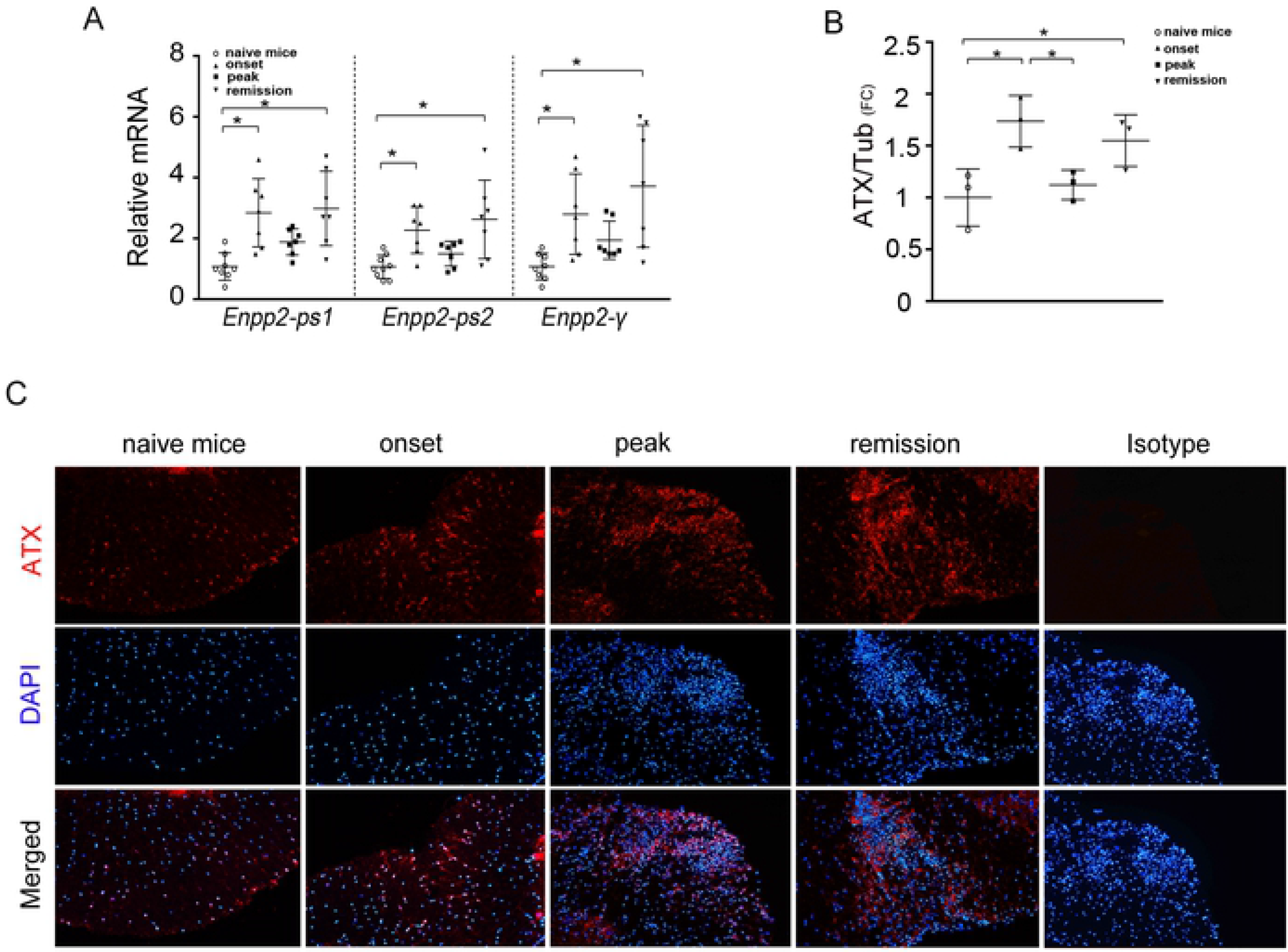
Increased ATX expression levels in the inflamed spinal cord upon EAE. (A) Q-RT-PCR analysis of total ATX (*Enpp2*) mRNA levels (with two different set of primers; ps1-2), as well as of the ATX brain-specific γ isoform in whole spinal cord lysates. (B) Densitometry analysis of ATX protein levels in spinal cords (shown in Figure S3). Expression was normalized to the levels of β-tubulin. (C) Representative immunofluorescent staining of spinal cords from mice sacrificed at different time points post immunization with an anti-ATX antibody (Cayman). All values are presented as mean (±SD); Statistical significance between experimental groups was assessed with one-way ANOVA complemented with Bonferroni or Dunn’s multiple pair test; *denotes statistical significance (p<0.05).

A similar ATX expression profile was obtained upon western blot analysis of spinal cord homogenates with a highly specific monoclonal antibody (Fig. 3B and S3A). Immunofluorescent staining of ATX in transverse spinal cord cryo-sections confirmed elevated ATX protein levels at the remission phase (but not the onset), and localized staining in inflamed white matter lesions during EAE progression (Fig. 3C); a similar staining profile was obtained with yet another antibody (Fig. S3B). Therefore, EAE development is accompanied by increased ATX levels in the spinal cord, peaking at the remission phase. However, it cannot be excluded that some ATX staining could be due to soluble ATX bound to the cell surface by integrins or other transmembrane or membrane associated molecules [28–30], whose expression could be modulated upon EAE.

Increased total LPA levels were also detected with HPLC/MS/MS in the spinal cord of mice at the remission phase of EAE (Fig. 4A), predominated, unlike the plasma, by the 18:2 species (Fig. S1A). The differences in total LPC levels in the spinal cord upon EAE did not reach statistical significance (Fig. 4B), while there was no correlation with the corresponding LPA species (Fig S1A). No significant changes in the mRNA profile of PLPPs, largely responsible for extracellular LPA degradation [31], were noted (Fig. 4C), suggesting minor involvement in the regulation of spinal cord LPA levels in EAE. Furthermore, the mRNA levels for the different receptors of LPA were found to fluctuate during EAE development (Fig. 4D), suggesting exacerbated LPA signaling in the inflamed spinal cord.

**Fig 4.**
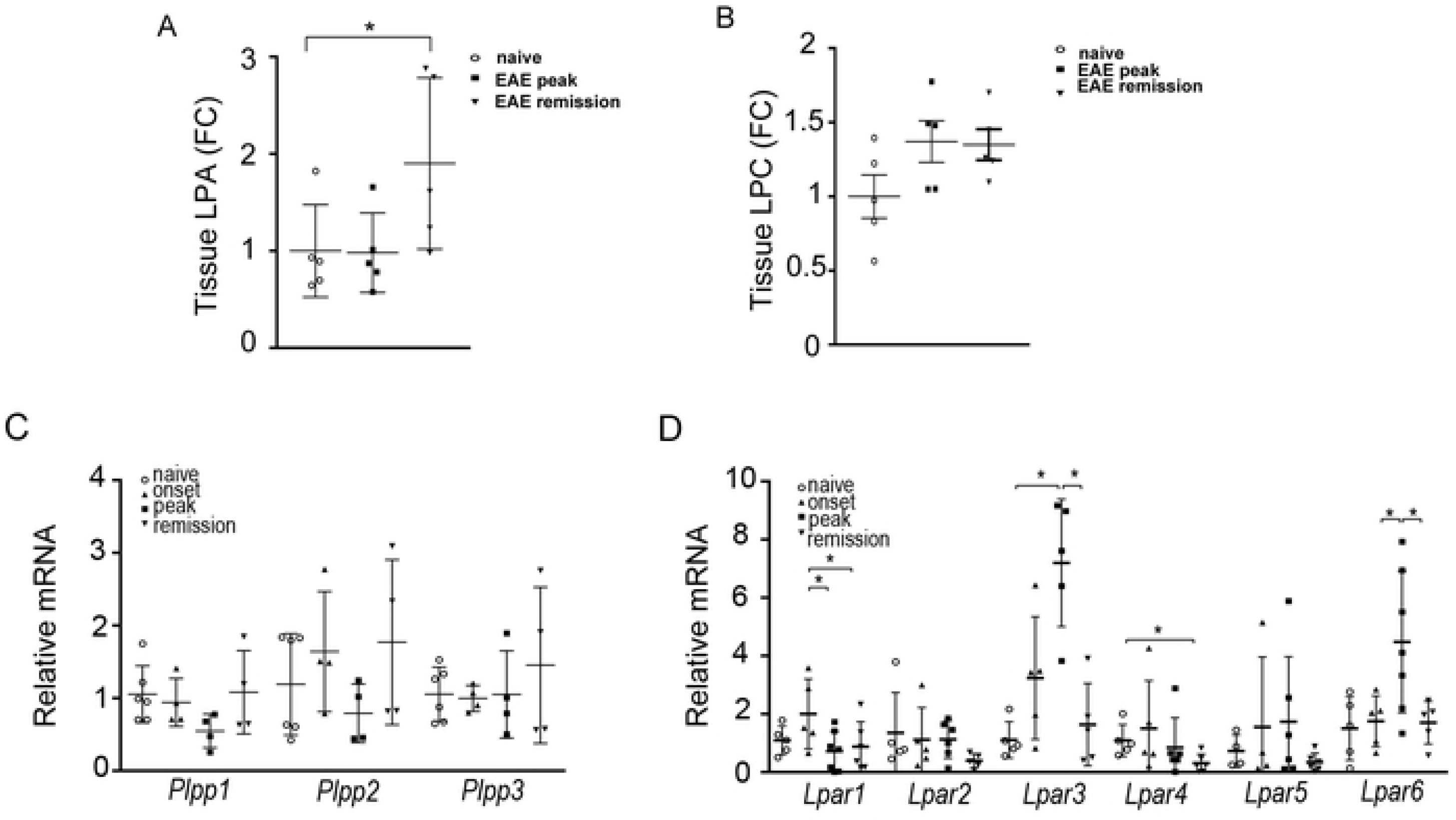
Deregulated LPA homeostasis and signaling in the spinal cord upon EAE. Total LPA (A) and LPC (B) levels in spinal cords; values were normalized to internal standards and presented as fold change (means ±SEM) to control samples. (C, D) Q-RTPCR analysis of spinal cords from mice undergoing EAE development for mRNA levels of (C) lipid phosphatases (*Plpps*) and (D) LPA receptors (*Lpars*) Values were normalized to the expression values of hypoxanthine-guanine phosphoribosyltransferase (HPRT) and are presented as mean fold change to controls (±SD). Statistical significance was assessed with one-way ANOVA complemented with Bonferroni or Dunn’s multiple pair test accordingly; *p<0.05.

Moreover, and given the suggested interplay of the PLA2/LPC and ATX/LPA axes in pathophysiology [32] we next examined the mRNA expression levels of different PLA2 isoforms. Increased mRNA levels of PLA2 g4a, g4c, g6, g7 and g15 were detected (Fig. S4A), suggesting *de novo* LPC production in the inflamed spinal cord that could stimulate ATX expression locally; PLA2-mediated LPA production cannot be excluded [7]. The mRNA levels of COX-1/2, guiding the synthesis of pro-inflammatory eicosanoids from PLA2-synthesized arachidonic acid (AA), were also found increased in EAE (Fig. S4B), indicating the parallel activation of multiple lipid pathways. Accordingly, a deregulation of various lipids was detected in the spinal cords of EAE mice, such as sphingomyelins and ceramides, as well as unsaturated fatty acids (Fig. S1B) suggesting an overall modulation of lipid metabolism upon EAE development and urging for further lipidomic studies, in both animal models as well as human patients.

Therefore, EAE development is accompanied with a deregulation of a series of molecules involved in lysophospholipid homeostasis and signaling including ATX, suggesting that the ATX/LPA axis has a possible role in EAE pathogenesis, likely in concert with the PLA2/LPC axis.

### ATX is expressed, among others, from activated macrophages and microglia during EAE pathogenesis

Given the important role of microglia and macrophages in EAE pathogenesis [4] and the suggested pathologic role of ATX expression from macrophages in modeled pulmonary inflammation and fibrosis [33], we next examined if inflammatory macrophages or resident microglia express ATX upon EAE development. Towards that end, FACS analysis of spinal cord homogenates was performed, utilizing the most widely accepted, but not exhaustive, FACS classification set [27]. The results (Fig. 5 A-B) indicate that upon EAE induction both microglial cells (CD45^low^CD11b^+^), as well as infiltrating blood-derived myeloid cells (CD45^hi^CD11b^+^) express ATX. Moreover, most macrophages and microglia expressing ATX upon EAE were positive for F4/80 expression (Fig. 5C), a well-known macrophage activation marker [4], suggesting that ATX/LPA could have an autocrine role in macrophage/microglia activation and/or maturation upon EAE.

**Fig 5.**
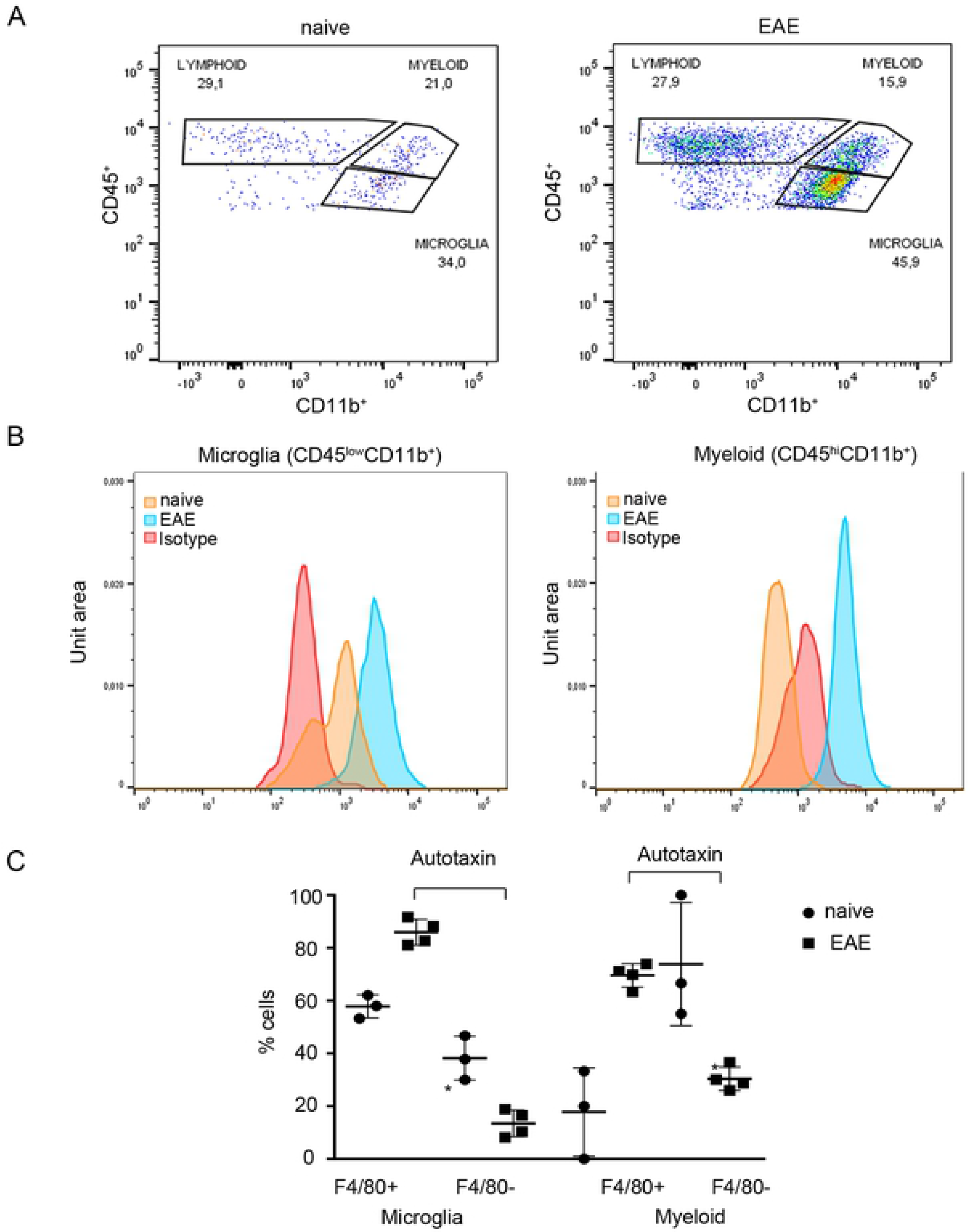
ATX is expressed by activated macrophages and microglia. Representative flow cytometry profile from spinal cord suspension prepared from naive and EAE mice. (A) CD45/CD11b FACs plot of EAE and naive mice following forward scatter and side scatter gating and doublets exclusion from CD45^+^ cells. lymphoid cells: CD45^+^CD11b^−^; microglia: CD45^low^CD11b^+^; myeloid cells: CD45^hi^CD11b^+^. (B) Representative ATX expression in microglia and macrophages in EAE and naive mice and isotype staining control. (C) F4/80 expression in ATX expressing CD11b^+^ cells. Statistical significance between experimental groups was assessed with one-way ANOVA complemented with Bonferroni multiple pair test. * p<0.05.

### ATX genetic deletion from CD11b^+^ cells decreases EAE severity

In order to explore a possible pathogenic role of ATX expression from macrophages/microglia during EAE, ATX was genetically deleted in these lineages by mating the conditional knockout mouse for ATX (*Enpp*2^f/f^)[9] with a transgenic mouse line expressing the Cre recombinase under the control of the CD11b promoter (Tg*CD11b-Cre*)[23]. *CD11bEnpp2*^−/−^ mice were born with a mendelian ratio and had properly recombined the *Enpp2* gene as qualitatively shown with PCR of DNA extracted from spinal cord sections (Fig. 6A).

**Fig 6.**
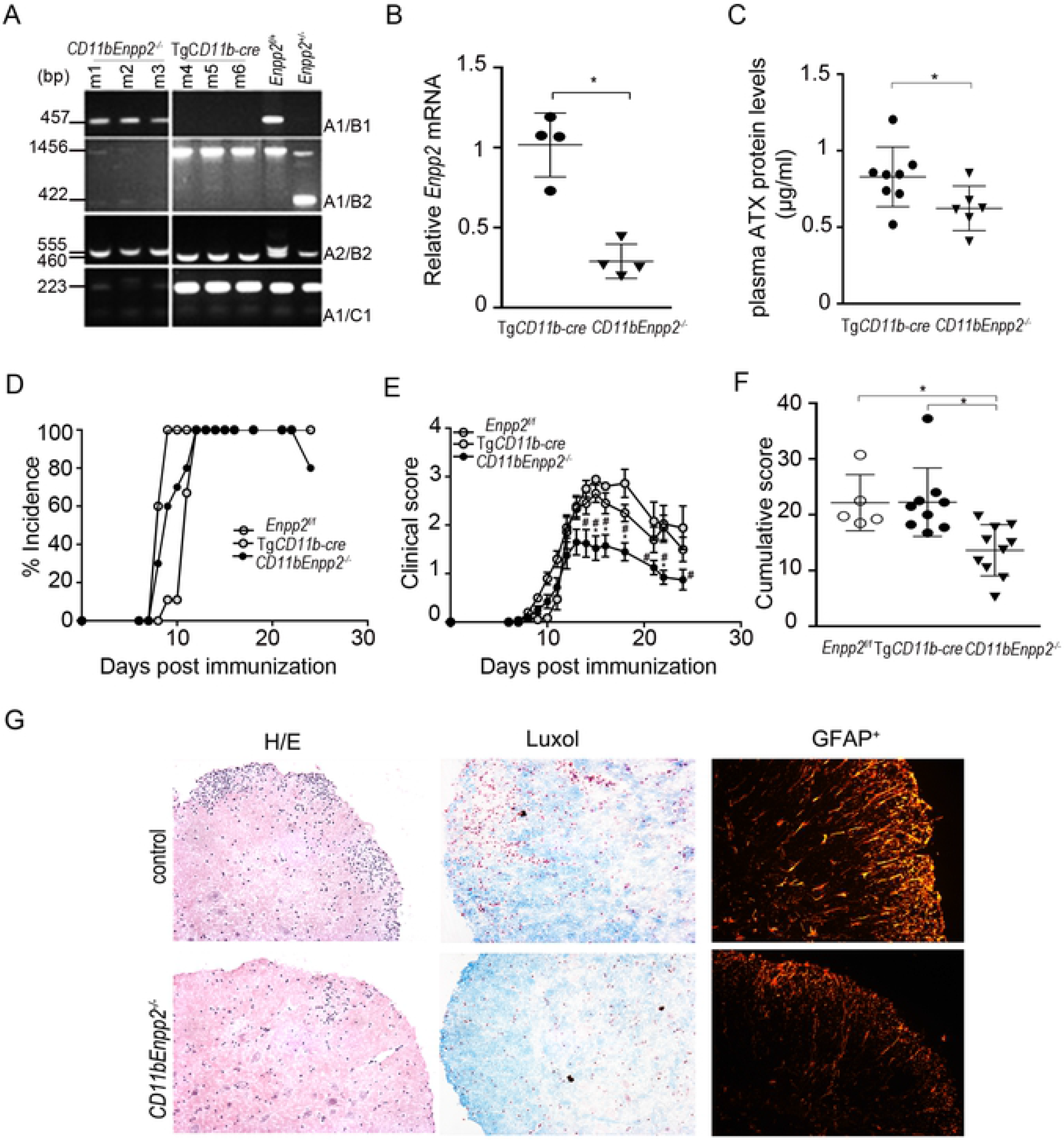
ATX genetic deletion from CD11b^+^ cells attenuates EAE severity. (A) Genomic PCR analysis indicates appropriate recombination in *CD11bEnpp2*^−/−^ mice. (B) Q-RT-PCR analysis of *Enpp2* mRNA levels in the spinal cords of *CD11bEnpp2*^−/−^ mice and control littermates. (C) Plasma ATX protein levels. (D) EAE incidence, (E) clinical scores, and (F) cumulative scores of EAE progression in *CD11bEnpp2*^−/−^ mice and littermate controls. (G) Representative H/E and Luxol fast blue stained spinal cord sections and immunofluorescent staining for astrocytes (GFAP^+^ cells). Values are presented as mean (± SD). Statistical significance between experimental groups was assessed with Mann-Whitney Rank Sum Test (E), one-way ANOVA complemented with Bonferroni multiple pair test (F) or Student t-test (B and C); *denotes statistical significance (p<0.05).

Genetic deletion of ATX from monocytic cells, decreased mRNA levels in the spinal cord (Fig. 6B), as well as ATX activity levels in the plasma (Fig. 6C), confirming both ATX expression from CD11b^+^ cells, as well as efficient genetic targeting. *CD11bEnpp2*^−/−^ mice did not exhibit any behavioral phenotype, nor any appreciable effect in CNS gross morphology under healthy conditions; no major effect in EAE incidence (Fig. 6D) or onset (Fig. 6E) was observed either. However, EAE progression and severity in *CD11bEnpp2*^−/−^ mice was significantly attenuated, with mice presenting with less disability at the peak of disease (Fig. 6E), resulting in a decreased cumulative score (Fig. 6F); no effect of the conditional *Enpp2* allele or the transgenic expression of the cre recombinase alone was observed in littermate controls in multiple experiments (Fig. 6 D-F). Histological analysis of *CD11bEnpp2*^−/−^ mice spinal cords indicated decreased inflammation, as detected with H/E staining, decreased demyelination, as detected with luxol staining, as well as significantly less activated astrocytes (GFAP^+^), an EAE hallmark (Fig. 6G).

Therefore, ATX expression from CD11b^+^ cells plays an important role in disease severity, likely including the LPA-mediated autocrine activation of macrophages, central to EAE pathogenesis.

## Discussion

In this report, increased ATX and LPA levels were found in the plasma and spinal cords of mice undergoing EAE development, amid an overall deregulated lipid homeostasis. More importantly, CD11b^+^ cells, mostly macrophages and microglia, were shown to express ATX upon EAE in the spinal cord, likely resulting to their autocrine activation. Genetic deletion of ATX from CD11b^+^ cells ameliorated the progression of EAE, thus proving an overall detrimental role of monocytic ATX expression and LPA signaling in EAE pathogenesis.

Increased ATX/LPA levels were detected in the plasma at the peak of EAE (Fig. 1 and S1), declining to normal levels during remission. Noteworthy, even further decreases of LPA were reported for the second remission phase (35 days post MOG immunization) in a SJL EAE model [21]. In MS patients, ATX/LPA plasma measurements have been conflicting [17–21], complicated by treatment history, the timing of sample collection and the available control samples; however, ATX/LPA serum/plasma levels were found higher during relapses comparing to remissions, in the largest and more recent studies [19, 21].

Given the suggested LPA effects in endothelial physiology [8, 32], the increased BBB permeability upon EAE development [2, 3], and the observations made here that ATX/LPA peak in the plasma earlier than in the spinal cord, it is tempting to assume that increased amounts of ATX/LPA could be extravasated in the CNS, possibly contributing to local LPA levels and promoting disease (EAE/MS) pathogenesis. However, and non-withstanding possible effects due to genetic modifications, no differences were observed between *Enpp2*^+/−^ or Tg*Enpp2*^+/+^ mice and their littermate controls in EAE pathogenesis (Fig. 2), suggesting that a systemic 2-fold (but life-long) fluctuation of ATX/LPA levels *per se* is likely not a disease modifying factor in the EAE model. Moreover, ATX has been suggested to modulate lymphocyte trafficking in lymphoid organs, where is highly expressed from the endothelial cells of high endothelial venules (HEV)[12, 34, 35]. However, possible ATX/LPA effects on lymphocyte trafficking, that could complement the well-recognized and therapeutically targeted Sphingosine-1-phosphate (S1P) effects (Brinkmann, Billich et al. 2010, Knowlden and Georas 2014), were not examined here.

Increased ATX, mRNA and protein, expression in the spinal cords of mice were detected during the development of EAE (Fig. 3). ATX levels were found significantly higher at the remitting phase of the disease (Fig. 3), rather than its peak as detected in the plasma. A similar mRNA expression profile was also found for the ATX γ isoform (Fig 3A), which is predominantly expressed in the CNS [13], suggesting that at least one (undetermined) part of the increased levels of ATX is derived locally from resident CNS cells. Little is known on the regulation of ATX expression in the CNS and especially in the context of EAE. TNF and IL-6, intricately linked with EAE development [24, 25], have been reported to stimulate ATX mRNA expression in different, non-CNS, cell types [6, 36, 37]; both TNF and IL-6 mRNA levels were found increased in EAE (Fig. S2A), suggesting that they could promote ATX expression in some cellular types, in a spatiotemporal manner.

LPC, the substrate of ATX, has been shown to directly promote ATX expression in hepatocytes [37], while it is well known to promote demyelination both *in vitro* and *in vivo* [32]. Although LPC is also a signaling lipid, many of the previously reported LPC effects, following the discovery of the lysophospholipase D properties of ATX, are now attributed to ATX and the conversion to LPA. Increased expression of different PLA2 isoforms, largely responsible for LPC synthesis (and possibly some LPA), were detected in the spinal cord upon EAE (Fig. S4A), although no significant changes were detected in spinal cord and plasma LPC levels (Fig. 4B, S1A). However, *de novo* synthesis could be masked by the high concentrations of LPC. Intriguingly, a search for putative functional partners of ATX at <u>STRING</u> returns mostly PLA2 isoforms, suggesting a physical association of ATX and PLA2s, possibly at the cell surface (via integrins and phosphatidylcholine respectively), and intriguing possibilities in the regulation of local LPC and LPA production and utilization during EAE pathogenesis, especially since integrin-bound-ATX-mediated LPA is thought to be engaged by the adjacent LPA receptors.

Despite the short life of LPA and its likely local consumption, that can complicate conclusions from LPA measurements, the LPA expression profile in the inflamed spinal cord followed the ATX one (Fig. 4), consistent with the consensus notion that ATX is largely responsible for the majority of extracellular LPA production [5–7]. No correlation was observed with plasma LPA both in terms of predominating species and as well as of timing, arguing against a major plasma contribution to spinal cord LPA levels; however, it cannot be excluded. A similar LPA spinal cord expression profile has been reported for the SJL EAE model [21], while increased LPA levels in the spinal cord have been also reported upon contusive injury [38], further supporting deregulated LPA homeostasis upon CNS damage.

Among the different potential cellular sources of ATX in the CNS, infiltrating myeloid cells, mainly macrophages, and resident microglial cells (CD11b^+^ cells) were found to express ATX (Fig. 5). In support, lung macrophages post bleomycin-mediated epithelial damage have been shown to produce ATX leading to LPA production in the bronchoalveolar fluid, and promoting the pathogenesis of pulmonary fibrosis [33]. LPS-mediated TLR activation of monocytic THP-1 cells was also reported to lead to ATX production [39]. Moreover, it was recently reported that LPA stimulates the expression of F4/80, a well-known macrophage activation marker [4], in bone marrow-derived and splenic monocytic (CD11b^+^) cells [40]. Accordingly, most macrophages and microglia (CD11b^+^ cells) expressing ATX upon EAE were positive for F4/80 expression (Fig. 5C), suggesting that the pathogenetic effect of ATX expression from monocytic cells in the context of EAE includes their activation. Accordingly, microglial activation has been shown to mediate *de novo* LPA production in a model of neuropathic pain [41], while intraspinal injection of LPA was suggested to induce macrophage/microglia activation [38].

Macrophage biology in the CNS remains understudied and controversial [4]. It is generally accepted that macrophages and microglia, that were both shown here to express ATX, can have differential roles in the pathogenesis of CNS disorders [4]. In EAE, myeloid cells are thought to have a greater role, promoting disease onset and progression. By contrast, microglia have been suggested to have a beneficial role, especially in the remitting phase, by aiding in tissue repair and remyelination [4]. The observed ATX expression from macrophages would stimulate their activation and effector functions, central to EAE pathogenesis, whereas ATX expression from microglia could promote wound healing and recovery from the disease. Although the overall relative contribution of macrophages and microglia, and the corresponding ATX expression, to EAE severity remains to be further explored, the observed EAE attenuation upon the genetic attenuation of ATX expression from CD11b^+^ cells (Fig. 6), can likely be attributed to the abolishment of its expression from inflammatory macrophages. However, further studies are required, especially since there are other types of CNS resident macrophages, namely perivascular, meningeal and choroid plexus macrophages [4].

Beyond the constitutive ATX expression from choroid plexus and leptomeningeal cells, and the overall pathogenic ATX expression from activated CD11b^+^ cells upon EAE, other cell types can potentially express ATX upon EAE and further modulate its pathogenesis. ATX has been shown to be expressed from oligodendrocyte precursor cells (OPCs) promoting their differentiation, via LPA and/or its matricellular properties [14, 42–44]. ATX expression from OPCs during EAE could promote their maturation and the increased production of oligodendrocytes, thus promoting remyelination, in support of a beneficial role for ATX/LPA in EAE pathogenesis. Moreover, activated astrocytes, that orchestrate CNS tissue repair following injury [45], have been suggested to express ATX upon neurotrama [15]. Multiple LPA effects to astrocyte physiology have been reported, including several of the hallmarks of reactive astrogliosis such as cytoskeletal re-organization, proliferation and axonal outgrowth [8]. Finally, and besides the possible ATX autocrine effects, multiple LPA effects in neuronal cell types have been reported [8], further supporting a multifaceted role for the ATX/LPA axis in EAE/MS pathophysiology. Importantly, the overall effect of ATX activity in EAE pathogenesis depends on the LPA spatiotemporal homeostasis, as well as the relative abundance of the different LPA receptors in the different CNS cell types. Noteworthy, anti-inflammatory roles have been suggested for the LPA receptor 2 on immune cells [46, 47], and *Lpar2* null mice were found protected from EAE development [21]. The identification of the specific LPA receptors for each cell type and their spatiotemporal activity will be crucial in understanding the multiple LPA cellular effects in the context of EAE pathogenesis.

The attenuation of EAE severity by the genetic deletion of ATX from CD11b^+^cells, also considering the other likely sources of ATX expression in the inflamed CNS, suggest possible therapeutic benefits of targeting ATX in EAE/MS. Indeed, pharmacologic potent ATX inhibition was recently reported to attenuate the development of EAE [48], timely with the booming discovery of novel ATX inhibitors [49].

## Conclusions

The ATX/LPA axis is increasingly recognized as a major player and drug target in different chronic inflammatory conditions and/or diseases. The present study reveals a novel detrimental role for macrophage (CD11b^+^) ATX expression in EAE development, urges further studies on LPA homeostasis in MS/EAE pathogenesis and supports potential therapeutic benefits from targeting ATX.

## Supporting information

**S1 Fig. Deregulated lipid metabolism upon EAE pathogenesis.** (A) Heat map of LPA and LPC species levels as measured with HPLC/MS/MS in plasma (n=10-12) and tissue (n=5) of mice upon EAE pathogenesis. (B) Heat map of sphingomyelins, ceramides and unsaturated fatty acid species measured with HPLC MS/MS in spinal cord tissue (n=5) of EAE mice compared with naïve mice.

**S2 Fig. Pro-inflammatory and pro-fibrotic mRNA expression in EAE.** Q-RTPCR analysis of spinal cords from mice undergoing EAE development for mRNA levels of (A) *Tnf-a* and *Il-6*, (B) *Tgf-b* and *Il-10* and (C) collagens (*col1a1, col3a1, col4a1*). Values were normalized to the expression values of hypoxanthine-guanine phosphoribosyltransferase (HPRT) and are presented as mean fold change to controls (±SD). Statistical significance was assessed with one-way ANOVA complemented with Bonferroni or Dunn’s multiple pair test accordingly; *p<0.05.

**S3 Fig. Increased ATX protein levels during EAE pathogenesis.** (A) Representative western blot (out of two) of ATX (and β-tubulin) expression in whole spinal cord lysates from mice undergoing EAE development, at the indicated time points. (B) Increased ATX protein levels during EAE pathogenesis in spinal cords. Representative (out of >5) immunofluorescent staining of spinal cords from mice sacrificed at different time points post immunization. 7μm sections were stained with an anti-ATX ab (Sigma) and the nuclei were counterstained with DAPI.

**S4 Fig. Increased *Pla2* and *Cox* mRNA expression in EAE**. Q-RTPCR analysis of spinal cords from mice undergoing EAE development for mRNA levels of **(A)** phospholipase A2 isoforms (*Pla2s*) and **(B)** *Cox-1/2*. Values were normalized to the expression values of hypoxanthine-guanine phosphoribosyltransferase (HPRT) and are presented as mean fold change to controls (±SD). Statistical significance was assessed with one-way ANOVA complemented with Bonferroni or Dunn’s multiple pair test accordingly; *p<0.05.

## Acknowledgments

The transgenic *TgEnpp2*^+/+^ mouse was kindly provided by G. Mills. We are thankful to M. Denis for and valuable advice on EAE and I. Barbayianni for assistance in the preparation and submission of the manuscript.

## Author Contributions

IN and IS performed most of the presented experiments. EK, GS and GP performed the MS/MS analyses. ST and CM performed Q-RT-PCR analyses. JA and GK provided expertise, antibodies and genetically modified mice. IS and VA designed the study and wrote the manuscript, which was critically read by all authors.

